# Transcriptional response of *Streptomyces coelicolor* to rapid chromosome relaxation or long-term supercoiling imbalance

**DOI:** 10.1101/602359

**Authors:** MJ Szafran, M Gongerowska, T Małecki, MA Elliot, D Jakimowicz

## Abstract

Negative DNA supercoiling allows chromosome condensation within a cell and facilitates DNA unwinding, which is required for the occurrence of DNA transaction processes, i.e., DNA replication, transcription and recombination. In bacteria, changes in chromosome supercoiling impact global gene expression; however, the limited studies on the global transcriptional response have focused mostly on pathogenic species and have reported various fractions of affected genes. Furthermore, the transcriptional response to long-term supercoiling imbalance is still poorly understood. Here, we address the transcriptional response to both novobiocin-induced rapid chromosome relaxation or long-term topological imbalance, both increased and decreased supercoiling, in environmental antibiotic-producing bacteria belonging to the *Streptomyces* genus. During the *Streptomyces* complex developmental cycle, multiple copies of GC-rich linear chromosomes present in hyphal cells undergo profound topological changes, from being loosely condensed in vegetative hyphae, to being highly compacted in spores. In *Streptomyces*, chromosomal supercoiling changes may also be triggered by environmental stressors and have been suggested to be associated with the control of antibiotic production. Remarkably, in *S. coelicolor*, one of model *Streptomyces* species, topoisomerase I (TopA) is solely responsible for the removal of negative DNA supercoils. Using a *S. coelicolor* strain in which *topA* transcription is under the control of an inducible promoter, we generated a long-term supercoiling imbalance, enabling us to identify genes involved in the supercoiling-sensitive transcriptional response. We observed that affected genes are preferentially organized in clusters, and among them, we identified a supercoiling-hypersensitive cluster (SHC) located in the core of the *S. coelicolor* chromosome. Moreover, using a gyrase inhibitor, we identified the directly affected novobiocin-sensitive genes and established that the AT content in their promoter regions was increased. Notably, genes whose expression was immediately impacted by gyrase inhibition encoded products associated with membrane transport or secondary metabolite synthesis. In contrast to the novobiocin-sensitive genes, the transcripts affected by long-term topological imbalance encompassed genes encoding nucleoid-associated proteins, DNA repair proteins and transcriptional regulators, including multiple developmental regulators. Collectively, our results show that long-term supercoiling imbalance globally regulates gene transcription and has the potential to impact development, secondary metabolism and DNA repair, amongst others.

## INTRODUCTION

A bacterial chromosome is highly constrained within the cell, yet remains fully accessible for DNA replication, segregation and transcription. In bacteria, these processes are not separated in space and time, and their co-occurrence significantly impacts chromosome architecture. Chromosome topology is sustained by the coordinated action of nucleoid-associated proteins (NAPs), condensins (SMC, MukB) and DNA topoisomerases (Bjorkegren and Baranello, 2018; Dame, 2005; Dillon and Dorman, 2010; Racko et al., 2018; Rovinskiy et al., 2012). Whereas NAPs are responsible for local DNA bending, wrapping and bridging, which organize the bacterial chromosome into topologically independent domains, condensins and topoisomerases induce global compaction and control overall chromosome negative supercoiling (Lindow et al., 2002; Postow et al., 2004). Moreover, topoisomerases are critical for the removal of excessive DNA supercoiling generated by DNA replication and transcription (Baaklini et al., 2008). All known bacterial genomes encode at least two essential topoisomerases – topoisomerase I (TopA and TopoI) and gyrase (GyrAB). Whereas TopA removes negative DNA supercoils, gyrase eliminates positive supercoils, thus, the opposite activities of TopA and GyrAB are essential for maintaining topological balance (topological homeostasis) (Champoux, 2001). Disturbances in the topological balance are reflected in the changes in transcriptional activity of the so-called supercoiling-sensitive genes (SSGs) (Peter et al., 2004). Decreased level of DNA supercoiling, resulting from gyrase inhibition, affects the transcription of approximately 7% of genes in *Escherichia coli* and up to 37% of genes in *Haemophilus influenzae*, (Gmuender et al., 2001; Peter et al., 2004). Similarly, increased supercoiling, generated by the inhibition of TopA activity, leads to changes in the transcription of 10% of all genes in *Streptococcus pneumoniae* (Ferrandiz et al., 2016). Although the importance of global DNA supercoiling on overall gene transcription in bacteria is well established, the scale of transcriptional changes differs between bacteria, and much is still not understood about these processes.

The genes most efficiently induced by the shift in DNA supercoiling are those that are critical for restoring topological balance, and encompass genes encoding topoisomerases (Ahmed et al., 2015; Szafran et al., 2016; Tse-Dinh, 1985), NAPs and condensins (Ghosh et al., 2014; Guha et al., 2018; Schneider et al., 2000). However, SSGs also include genes involved in the regulation of many other cellular processes, i.e., DNA replication, cell division, and metabolic pathways, as well as DNA repair or stress adaptation (Peter et al., 2004). Interestingly, even after the apparent restoration of supercoiling homeostasis, the transcription of many genes has been reported to remain affected (Ferrándiz et al., 2014). Since changes in chromosome supercoiling are induced by environmental stresses, including osmotic and oxidative stress, host invasion, and temperature down- or up-shift, transcriptional regulation by chromosome topology may be an important bacterial strategy for rapid adaptation to stress conditions (Camacho-Carranza et al., 1995; Cameron et al., 2011; Cheung et al., 2003; Horiuchi et al., 1984; Tse-Dinh et al., 1997; Weinstein-Fischer et al., 2000).

By virtue of their being exposed to a wide range of environmental stressors, soil bacteria have evolved a multitude of strategies to facilitate their survival in harsh conditions. A classic example is provided by the soil-dwelling *Streptomyces*, which are Gram-positive, mycelial bacteria that possess linear, GC-rich chromosomes (Bentley et al., 2002). *Streptomyces* environmental niche adaptations include a complex regulatory network that employs a high number of transcriptional regulators, including more than 60 sigma factors and approximately 60-70 of two component systems (TCSs). Moreover, *Streptomyces* possess a high number of DNA protection proteins and enzymes involved in the posttranslational modifications of proteins, i.e., proteases, acetyltransferases, kinases (de Crecy-Lagard et al., 1999; Facey et al., 2009; Hutchings et al., 2004). Finally, *Streptomyces* produce an assortment of secondary metabolites with varied biological activities (Jeong et al., 2016) and exhibit a complex life cycle that involves the formation of stress-resistant spores. Vegetatively growing *Streptomyces* develop branched multicellular hyphae composed of elongated cells containing multiple loosely-condensed chromosomes (Kim et al., 2000; McCormick and Flardh, 2012). During sporulation, the multigenomic unbranched aerial hyphae are converted into chains of unigenomic spores. This transition involves the condensation and segregation of tens of chromosomes, which in turn is coordinated with multiple synchronized cell divisions (Jakimowicz et al., 2005; Jakimowicz and van Wezel, 2012; Kois et al., 2009; Szafran et al., 2013). The different steps of *Streptomyces* differentiation are tightly regulated by a cascade of regulatory proteins (encoded by the *bld* and *whi* genes) that controls the expression of those genes involved in cell division and spore maturation (Bush et al., 2013; Flardh and Buttner, 2009; McCormick and Flardh, 2012).

*Streptomyces* sporulation involves prominent changes in chromosome organization and requires the activity of proteins that exert chromosome rearrangements, including topoisomerase TopA (Szafran et al., 2013). In contrast to other bacterial species, TopA is the only type I topoisomerase in *Streptomyces*. As in other actinobacteria, the enzyme differs remarkably from its bacterial homologs in having an unusually high processivity, which is provided by a long stretch of alanine/lysine repeats in its C-terminal domain (Bhaduri et al., 1998; Strzalka et al., 2017; Szafran et al., 2014). In *S. coelicolor* and *S. venezuelae*, two model species broadly used in the studies of *Streptomyces* biology, TopA depletion increased DNA supercoiling and lead to growth retardation as well as impaired spore production (Donczew et al., 2016; Szafran et al., 2013). Thus, in *Streptomyces*, the removal of the topological tensions by the activity of the highly processive and indispensable TopA is required for efficient growth and sporulation.

In *Streptomyces*, as in other bacteria (*e.g. E. coli, Mycobacterium smegmatis* and *S. pneumoniae*), novobiocin-mediated gyrase inhibition leads to the rapid loss of chromosome supercoiling (Ferrandiz et al., 2010; Guha et al., 2018; Peter et al., 2004; Szafran et al., 2016). In most bacteria, topological balance is subsequently restored by simultaneous upregulation of gyrase genes, and downregulation of *topA* transcription (Ferrandiz et al., 2010; Tse-Dinh, 1985; Unniraman and Nagaraja, 1999), while uniquely in *Streptomyces*, chromosome relaxation appears to affect only the *gyrBA* operon but does not influence *topA* transcription (Szafran et al., 2016). Conversely, increased negative chromosome supercoiling caused by depletion of TopA leads to the induction of *topA* expression, but, interestingly, does not influence *gyrBA* transcription. Changes in topoisomerase levels or superhelical density have also been observed as a result of *Streptomyces* exposure to increased temperature or osmotic stress (Aldridge et al., 2013; Szafran et al., 2016). Thus, *Streptomyces* provide an interesting model for studies of topological homeostasis and its role in stress responses, given their soil habitat and unique system of chromosome supercoiling maintenance.

Here, we analyzed the transcriptional response of *S. coelicolor* to changes in global chromosome supercoiling by inhibiting DNA gyrase activity. We identified those genes that were immediately impacted by the rapid loss of DNA supercoiling. Additionally, using the strain in which TopA level could be modified we defined the effect of constitutive and long-term topological imbalance on global gene expression.

## MATERIALS AND METHODS

### Bacterial strains

The *S. coelicolor* strains used in this study are listed in Table 1.

**Table 1.** Strains used in this study.

| Strain | Relevant genotype | Source or reference |
| --- | --- | --- |
| **M145** | SCP1<sup>-</sup> SCP2<sup>-</sup> | (Bentley et al., 2002) |
| **PS04** | M145 $\Delta topA::scar$ attBΦC31::pIJ6902$topA$ | (Szafran et al., 2013) |
| **PS08** | M145 pIJ6902 | (this study) |
| **MS10** | M145 pWHM3Hyg | (Szafran et al., 2016) |
| **MS11** | PS04 pWHM3Hyg | (Szafran et al., 2016) |

To construct the PS08 stain, wild type *S. coelicolor* (M145) was transformed with the integrative pIJ6902 plasmid (Huang et al., 2005) according to the procedures described by Kieser et al. (Kieser et al., 2000) and subsequently selected for apramycin resistance.

### Western blot analysis

To determine TopA protein levels, crude cell extracts were prepared from 18-hour cultures in 79 medium (Prauser and Falta, 1968). Cell lysate proteins (5 μg in total) were separated in a 10% denaturing polyacrylamide gel before being transferred to a nitrocellulose membrane. The TopA protein was subsequently detected using primary rabbit polyclonal anti-TopA antibodies and mouse secondary anti-rabbit IgG antibodies conjugated with alkaline phosphatase (Sigma Aldrich), according to the procedure described previously (Szafran et al., 2013). The relative protein level (RPL) was calculated using Fiji Software as the TopA band intensity compared to the wild type strain.

### Reporter plasmid isolation

The pWHM3Hyg plasmid, which served as a probe of the DNA supercoiling state *in vivo*, was isolated, according to the procedure described previously (Szafran et al., 2016), from *S. coelicolor* liquid cultures cultivated for 18 hours in YEME/TSB supplemented with 50 μg/ml hygromycin B (Kieser et al., 2000) at 30°C. To inhibit gyrase activity and induce rapid chromosome relaxation, novobiocin was added to the *S. coelicolor* culture to a final concentration of 10 μg/ml (approximately 1 × MIC_90_, which is the minimal concentration that inhibits *S. coelicolor* growth by 90%; Fig. S1) 10 minutes before mycelia were collected. The isolated plasmid DNA was resolved in a 0.8% agarose gel. To visualize particular topoisomers, the gel was stained with ethidium bromide. The topoisomer distribution was analyzed using ImageJ software.

### RNA isolation and RNA-Seq analyses

To follow the effects of altered DNA topology on cellular transcription, RNA was isolated from *S. coelicolor* mycelia cultivated in 30 mL YEME/TSB liquid medium for 18 hours. Mycelia were collected by centrifugation, frozen and stored at −70°C for subsequent RNA isolation. RNA was isolated using the procedure described previously by Moody et al. (Moody et al., 2013), digested with TURBO DNase I (Invitrogen, USA) and checked for chromosomal DNA contamination using PCR. The construction of strand-specific cDNA libraries, with an average fragment size of 250 bp, and sequencing using the MiSeq kit (Illumina) were performed by the Farncombe Metagenomics Facility at McMaster University (Canada). Paired-end 76-bp reads were subsequently mapped against the *S. coelicolor* chromosome using Rockhopper software (McClure et al., 2013), which enabled the successful alignment of 1.0-1.5 *10^6^ reads per sample (Fig. S2). For data visualization, Integrated Genome Viewer (IGV) software was used. The analysis of differentially regulated genes was based on the data generated by Rockhopper software. To calculate the fold change in gene transcription, normalized gene expression (normalization by the upper quartile of gene expression) of the wild type strain under control conditions was divided by normalized gene expression under particular experimental conditions. Subsequently, the log2 value of the fold change was calculated. Thetranscripts with a log2 value between −1.5 and 1.5 were eliminated from subsequent analyses, with the exception of NAPs and topoisomerases encoding genes, for which the threshold was based only on the p-value significance. The gene distribution analysis was performed by mapping the positions of transcription start points of identified genes or the first gene in a regulated operon determined on the basis of the RNA-seq data, to the *S. coelicolor* chromosome. To identify gene clusters of interest, we determined the percent of affected genes within a 250 kbp fragment of the *S. coelicolor* chromosome, subsequently moving the calculation window by 125 kbp in every step. Those regions in which the percentage of affected genes was higher than 5% were classified as supercoiling-sensitive clusters.

### Reverse-transcription and quantitative PCR (RT-qPCR)

For RT-qPCR analyses, RNA from *S. coelicolor* mycelia cultivated in liquid 79 medium for 24 hours was isolated using the GeneJET RNA isolation kit (Thermo Scientific, USA) according to the manufacturer’s procedure (except that the concentration of the lysozyme in the suspension buffer was increased to 10 mg/ml). The isolated RNA was digested with TURBO DNase I (Invitrogen) to remove traces of chromosomal DNA and then purified and concentrated using the GeneJET RNA Cleanup kit (Thermo Scientific). A total of 500 ng of RNA was used for cDNA synthesis using a Maxima First Strand cDNA synthesis kit (Thermo Scientific, USA) in a final volume of 20 μl. The original manufacturer protocol was modified for GC-rich *S. coelicolor* transcripts by increasing the temperature for first-strand synthesis to 65°C and extending the synthesis time to 30 minutes. Subsequently, the resulting cDNA was diluted to 100 μL and directly used for quantitative PCRs performed with PowerUp SYBR Green Master Mix (Applied Biosystems, USA). The relative level of a particular transcript was quantified using the comparative ΔΔCt method and the *hrdB* gene as the endogenous control (StepOne Plus real-time PCR system, Applied Biosystems, USA). The sequences of optimized oligonucleotides used in this study were synthetized by Sigma Aldrich (USA), and are listed in the supplementary data (see the supplementary file ‘*oligonucleotides*’).

### Analysis of the promoter AT content

To estimate the AT content in the promoter regions of novobiocin-induced genes, 66 DNA sequences (960 bp in length) encompassing 750 bp upstream of the translation start codons and 210 bp downstream were analyzed. As the control experiments, 66 randomly chosen promoter regions or 66 random *S. coelicolor* genomic sequences (each 960 bp in length) were used. Subsequently, we calculated the AT percentage of these sequences in 100-bp windows, with a 1-bp step. The AT content for each window was subsequently plotted against the first nucleotide position of the particular window. The differences of means calculated for specific nucleotide positions (340 bp and 660 bp) were tested using the Shapiro-Wilk model and T-test, respectively.

## RESULTS and DISCUSSION

### Rapid loss of chromosome supercoiling positively regulates *S. coelicolor* gene transcription

In previous work, we found that exposing *S. coelicolor* to novobiocin, initially led to the rapid relaxation of DNA, followed by gradual restoration of topological homeostasis (Szafran et al., 2016), as had been seen in other bacteria (Ferrandiz et al., 2010). The return to topological balance appeared to stem from changes in the expression of topoisomerase encoding genes, which in *S. coelicolor* corresponded to a strong induction of *gyrAB* but only slight inhibition of *topA* (Szafran et al., 2016); this is different from the simultaneous induction of *gyrAB* and inhibition of *topA* transcription observed in other bacteria (Ahmed et al., 2015; Unniraman and Nagaraja, 1999). Here, we sought to identify other genes in *S. coelicolor* whose transcription was also directly affected by the rapid chromosome relaxation. To this end, we treated *S. coelicolor* liquid cultures with 10 μg/ml novobiocin (the concentration equal to MIC_90_, Fig. S1). We analyzed the changes in global transcript abundance after 10 minutes of novobiocin exposure, and compared these with an untreated control strain. Based on previous work (Gmuender et al., 2001; Szafran et al., 2016), 10 minutes of novobiocin exposure was presumed to be sufficient to affect promoters directly sensitive to gyrase inhibition, triggering the primary transcriptional response, but was deemed insufficient time to induce a secondary transcriptional response (which depends on any primary-induced transcriptional regulators).

In analyzing our RNA-Seq data, we found that the rapid chromosome relaxation caused by novobiocin (Fig. 1A and S3A-B) led to a 2.71-fold induction of the *gyrBA* operon transcription (Fig. 1B), confirming that the primary transcriptional response could be induced by 10 minutes of gyrase inhibition. We next set out to identify genes exhibiting distinct changes in transcription in comparison to the untreated control (at least 2.83-fold), and found 121 genes that were sensitive to the rapid loss of chromosome supercoiling. These genes constitute 1.5% of the *S. coelicolor* open reading frames (ORFs) (see supplementary file ‘*novobiocin*’ for the complete list of the identified genes). Mapping of the transcription start sites (TSSs) of the identified genes to the *S. coelicolor* chromosome revealed that the novobiocin-sensitive genes were unevenly distributed along the *S. coelicolor* chromosome (Fig. 1C and S3C). Surprisingly, in contrast to *E*. coli, in which 106 genes were induced whereas 200 genes were repressed by gyrase inhibition (Peter et al., 2004), in *S. coelicolor* most of the novobiocin-sensitive genes (117 of 121) were upregulated by chromosome relaxation, with only 4 genes (*sco0498*, *sco1377, sco3726* and *sco3918*) being strongly downregulated (log2-fold>1.5).

**Figure 1.**
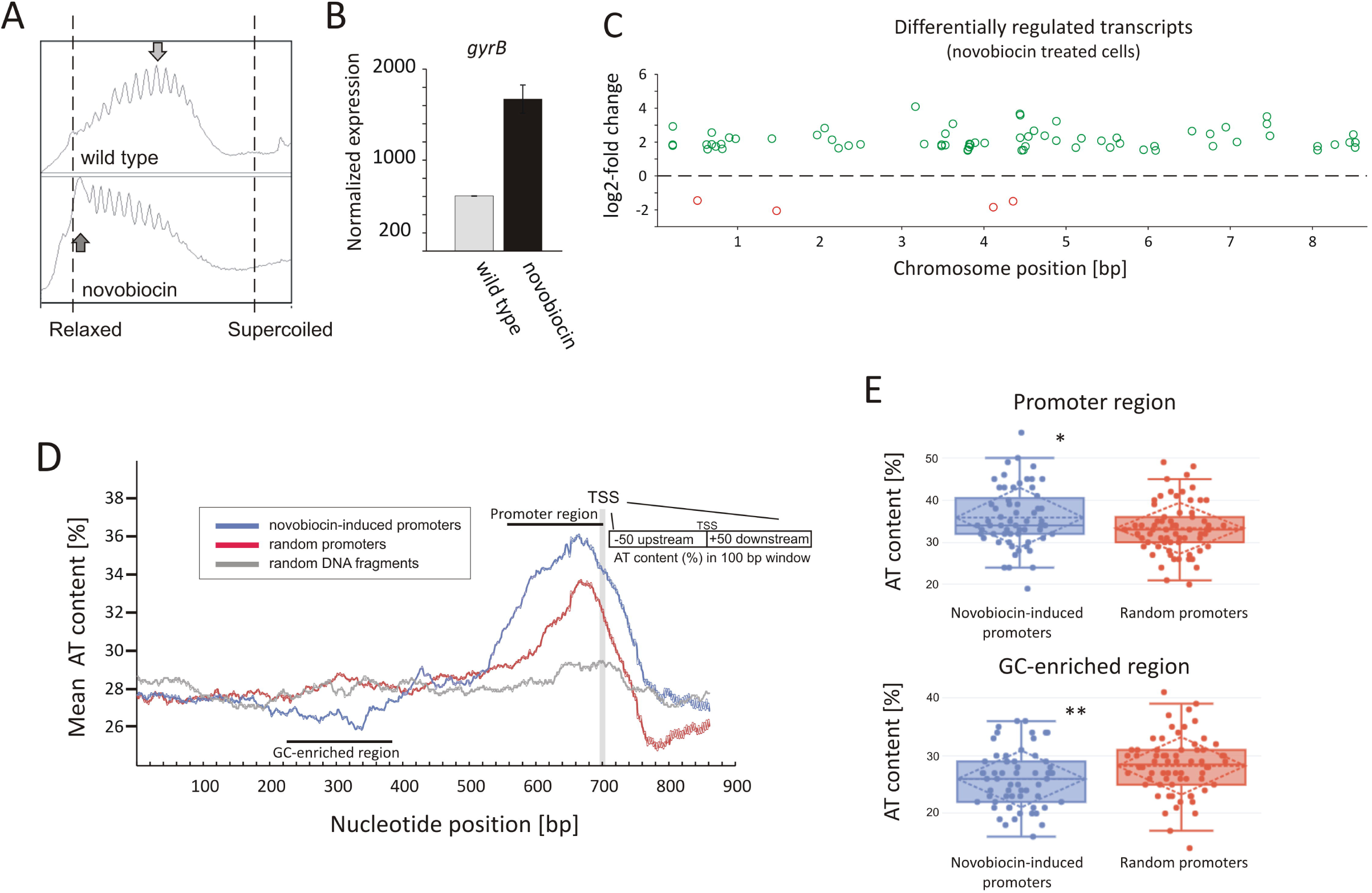
Identification of novobiocin-sensitive genes in *S. coelicolor*. **(A)** Analysis of reporter plasmid supercoiling density. The reporter plasmid was isolated from untreated wild type strain (upper panel) and the same strain exposed to novobiocin (10 μg/mL) for 10 minutes (bottom panel). The most abundant topoisomers identified under each condition are marked with arrows. **(B)** RNA-Seq based *gyrB* expression normalized by the upper quartile, in the untreated wild type strain (gray) and the novobiocin-treated strain (as described for (A)) (black). **(C)** The chromosomal localization of the novobiocin-sensitive genes. Differentially upregulated (green) and downregulated (red) transcripts in the *S. coelicolor* culture following novobiocin exposure (as in A) were identified using log2-fold changes>1.5; p value <0.05 thresholds. **(D)** Analysis of the average AT content in the promoter regions of novobiocin-sensitive genes (blue) in comparison to the random promoters (red) and random chromosomal sequences (gray). The transcriptional start site (TSS), promoter region and GC-enriched region identified upstream of novobiocin-sensitive promoters are marked. **(E)** The box plot comparison of AT content within the promoter region (upper panel) and GC-enriched region (bottom panel) of novobiocin-sensitive (blue) and randomly selected (red) promoters. The asterisk indicate the calculated p value <0.05 (*) or <0.01 (**).

We speculate that the gene upregulation after novobiocin exposure rather than their transcriptional repression may be due to the high GC content (72%) of the *S. coelicolor* genome (Bentley et al., 2002). In *E. coli*, promoter regions of relaxation-induced genes are notably enriched in AT base pairs, whereas relaxation-repressed promoters have increased GC content (Peter et al., 2004). In *S. coelicolor*, the increase in GC content in the promoter regions to levels above the average would be expected to result in increased DNA stability, which in turn may inhibit promoter unwinding. Conversely, more AT-rich promoter regions would be expected to favor promoter upregulation. To test this hypothesis, we compared the AT content of 66 novobiocin-induced promoters with the AT content of randomly chosen promoter regions. As predicted, we found that novobiocin-induced genes showed approximately 2-3% higher AT content compared with the randomly selected promoters and 8-9% higher in comparison to the average of the *S. coelicolor* genome (28%). Notably, the AT-rich region was significantly expanded in the novobiocin-sensitive genes relative to the random promoters (Fig. 1D). Unexpectedly, AT-content analysis revealed a GC-rich region located approximately 100-200 bp upstream of the novobiocin-induced genes (Fig. 1D), and not of the random promoters. The difference in the mean AT-content values between the novobiocin-sensitive and random promoters, calculated for position 340 bp (GC-enriched region) and 660 bp (promoter region) were statistically relevant (p values of 0.006 and 0.017, respectively; Fig. 1E). Interestingly, similar analysis performed in *E. coli* did not indicate the existence of GC-rich fragment preceding the relaxation-induced genes (Peter et al., 2004). Seemingly, the AT content within the promoters and its influence on transcriptional regulation may vary in dependence on overall chromosome organization. Although the role of the GC-enriched region upstream of the *Streptomyces* novobiocin-inducible promoters is unclear, we speculate that it may act as a DNA barrier for supercoil diffusion and/or may influence DNA curvature, thus affecting promoter susceptibility to unwinding.

Among the 121 novobiocin-sensitive genes, 15 encoded putative regulatory proteins. Included amongst these were the anti-sigma factor RsfA (Homerova et al., 2000), the developmental regulator WblC (Fowler-Goldsworthy et al., 2011), and the secondary metabolism regulator NsdB (Zhang et al., 2007). Interestingly, a high fraction of the novobiocin-sensitive genes (∼30%) encode proteins involved in either antibiotic production (10 genes, including the actinorhodin activator *sco5085/actII-orf4* and genes within the coelibactin biosynthetic cluster *sco7681-sco7688*) or putative membrane transporters (at least 25 genes) (Table 2). This suggests that rapid chromosome relaxation functions as the stress signal that triggers the transcription of genes whose products are involved in stress adaptation (i.e., those encoding regulatory proteins) and/or interspecies competition. Considering that decrease of gyrase activity and rapid chromosome relaxation may result from exposure to different cell membrane destabilizing agents that affect transmembrane potential and ATP synthesis, or other biologically active molecules produced by competing bacterial species (Prakash et al., 2009), it is conceivable that the intrinsic response to such agents should encompass the induction of transport proteins that may restore homeostasis to the intracellular environment and/or function to eliminate toxic compounds.

**Table 2.**
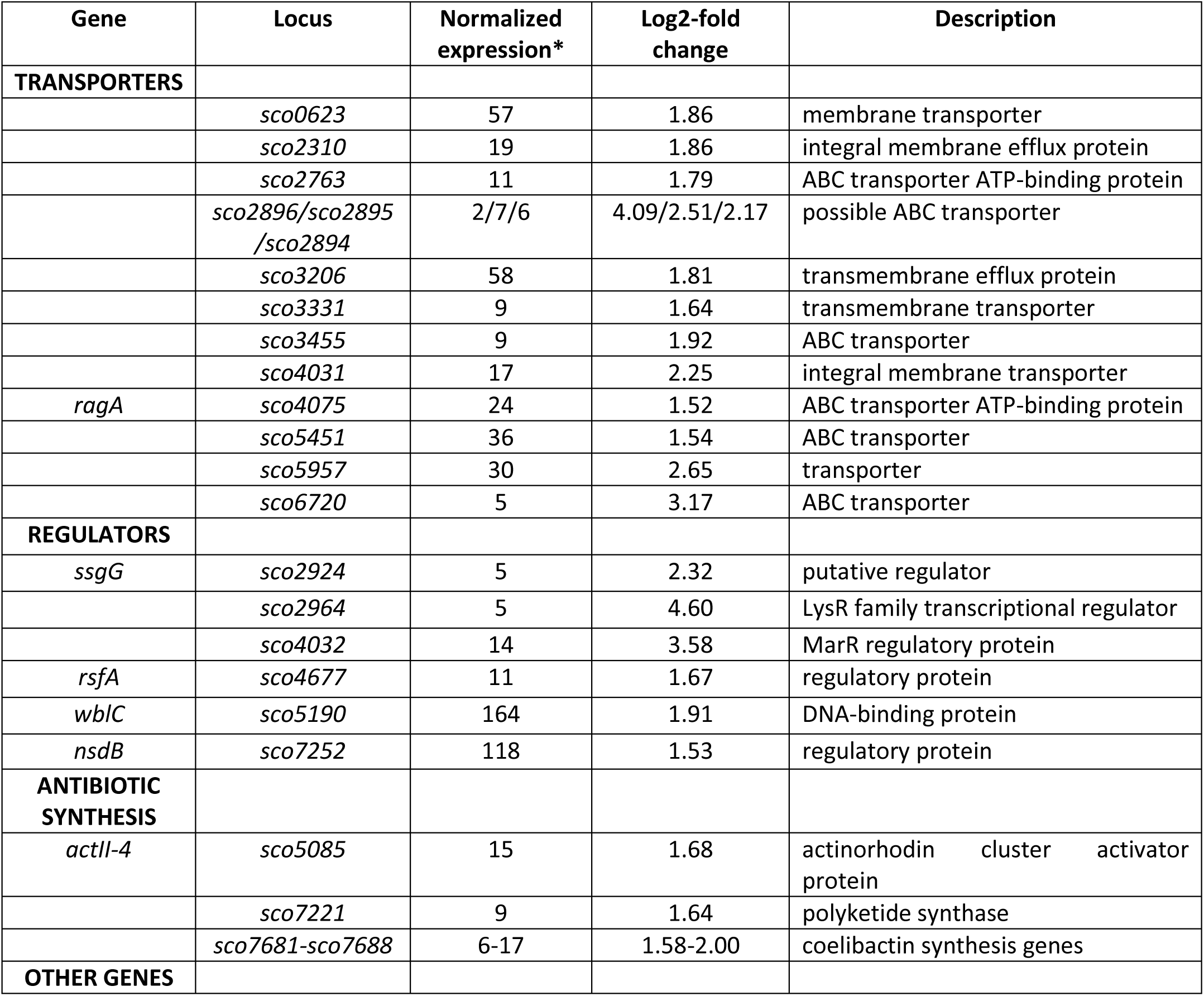
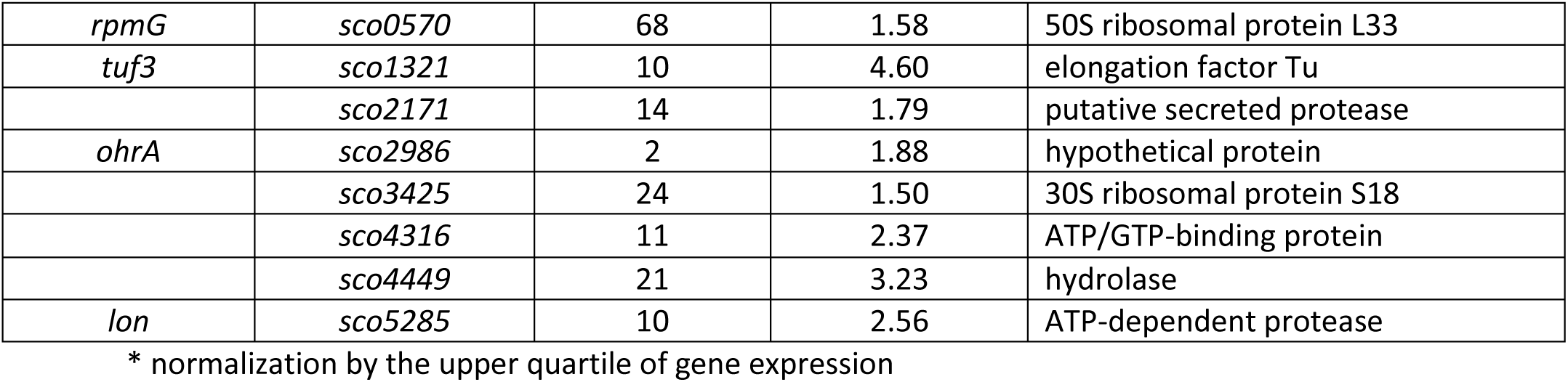
Selected novobiocin-induced genes encoding transporters or transcriptional regulators (log2-fold >1.5)

| Gene | Locus | Normalized expression* | Log2-fold change | Description |
| --- | --- | --- | --- | --- |
| **TRANSPORTERS** |  |  |  |  |
|  | *sco0623* | 57 | 1.86 | membrane transporter |
|  | *sco2310* | 19 | 1.86 | integral membrane efflux protein |
|  | *sco2763* | 11 | 1.79 | ABC transporter ATP-binding protein |
|  | *sco2896/sco2895/sco2894* | 2/7/6 | 4.09/2.51/2.17 | possible ABC transporter |
|  | *sco3206* | 58 | 1.81 | transmembrane efflux protein |
|  | *sco3331* | 9 | 1.64 | transmembrane transporter |
|  | *sco3455* | 9 | 1.92 | ABC transporter |
|  | *sco4031* | 17 | 2.25 | integral membrane transporter |
| *ragA* | *sco4075* | 24 | 1.52 | ABC transporter ATP-binding protein |
|  | *sco5451* | 36 | 1.54 | ABC transporter |
|  | *sco5957* | 30 | 2.65 | transporter |
|  | *sco6720* | 5 | 3.17 | ABC transporter |
| **REGULATORS** |  |  |  |  |
| *ssgG* | *sco2924* | 5 | 2.32 | putative regulator |
|  | *sco2964* | 5 | 4.60 | LysR family transcriptional regulator |
|  | *sco4032* | 14 | 3.58 | MarR regulatory protein |
| *rsfA* | *sco4677* | 11 | 1.67 | regulatory protein |
| *wblC* | *sco5190* | 164 | 1.91 | DNA-binding protein |
| *nsdB* | *sco7252* | 118 | 1.53 | regulatory protein |
| **ANTIBIOTIC SYNTHESIS** |  |  |  |  |
| *actII-4* | *sco5085* | 15 | 1.68 | actinorhodin cluster activator protein |
|  | *sco7221* | 9 | 1.64 | polyketide synthase |
|  | *sco7681-sco7688* | 6-17 | 1.58-2.00 | coelibactin synthesis genes |
| **OTHER GENES** |  |  |  |  |
| *rpmG* | *sco0570* | 68 | 1.58 | 50S ribosomal protein L33 |
| *tuf3* | *sco1321* | 10 | 4.60 | elongation factor Tu |
|  | *sco2171* | 14 | 1.79 | putative secreted protease |
| *ohrA* | *sco2986* | 2 | 1.88 | hypothetical protein |
|  | *sco3425* | 24 | 1.50 | 30S ribosomal protein S18 |
|  | *sco4316* | 11 | 2.37 | ATP/GTP-binding protein |
|  | *sco4449* | 21 | 3.23 | hydrolase |
| *lon* | *sco5285* | 10 | 2.56 | ATP-dependent protease |

\* normalization by the upper quartile of gene expression

In summary, the significant fraction of novobiocin-sensitive genes encompassed those encoding membrane transporters, antibiotic synthesis and regulatory proteins. The novobiocin-sensitive genes were nonuniformly distributed along the chromosome, and most of them were upregulated. The upregulation was reflected by the increased AT-content of their promoter regions and the presence of a upstream GC-rich DNA stretch.

### Long-term exposure to topological imbalance globally affects gene transcription

Having established that approximately 120 genes were directly impacted by gyrase inhibition, we were next interested in assessing the response of *S. coelicolor* to the long-term effects of altered chromosome supercoiling. To date, there is little understood about the long term effects of altered supercoiling, since most of studied bacteria were able to quickly restore topological homeostasis. Moreover, the effects of increased negative supercoiling on global gene transcription in bacteria remain relatively unexplored, in part due to the limited availability of selective TopA inhibitors (Cheng et al., 2013; Garcia et al., 2011; Godbole et al., 2014; Szafran et al., 2018).

To investigate long terms effects of supercoiling imbalance, we took advantage of a *S. coelicolor* strain (PS04) that had been engineered such that chromosome supercoiling could be modulated by tuning TopA levels using a thiostrepton-inducible *topA* gene (Szafran et al., 2013). In this PS04 strain, TopA depletion (no induction) dramatically increased the overall negative DNA supercoiling (Fig. 2A and S3A-B), but importantly did not stimulate *topA* expression, thus impeding the restoration of supercoiling balance. Moreover, inefficient repression of the *S. coelicolor* gyrase operon eliminated another possible pathway to restoring chromosomal topological balance in the TopA-depleted strain. On the other hand induction of *topA* expression in the PS04 strain led only to a slight (approximately 22%) increase of TopA protein level which also corresponded to a modest shift in global DNA supercoiling levels if compared to the wild type strain (Fig. 2A and S3A,B). This observation stays in agreement with our previous data showing that by increasing the concentration of *topA* inducer, only 25% increase of TopA level could be achieved (Szafran et al., 2013). This may be partially explained by induction of the *gyrBA* operon, which could compensate for high levels of TopA, and restore the supercoiling balance (Szafran et al., 2016) or by posttranscriptional regulation mechanisms of *topA* gene expression.

**Figure 2.**
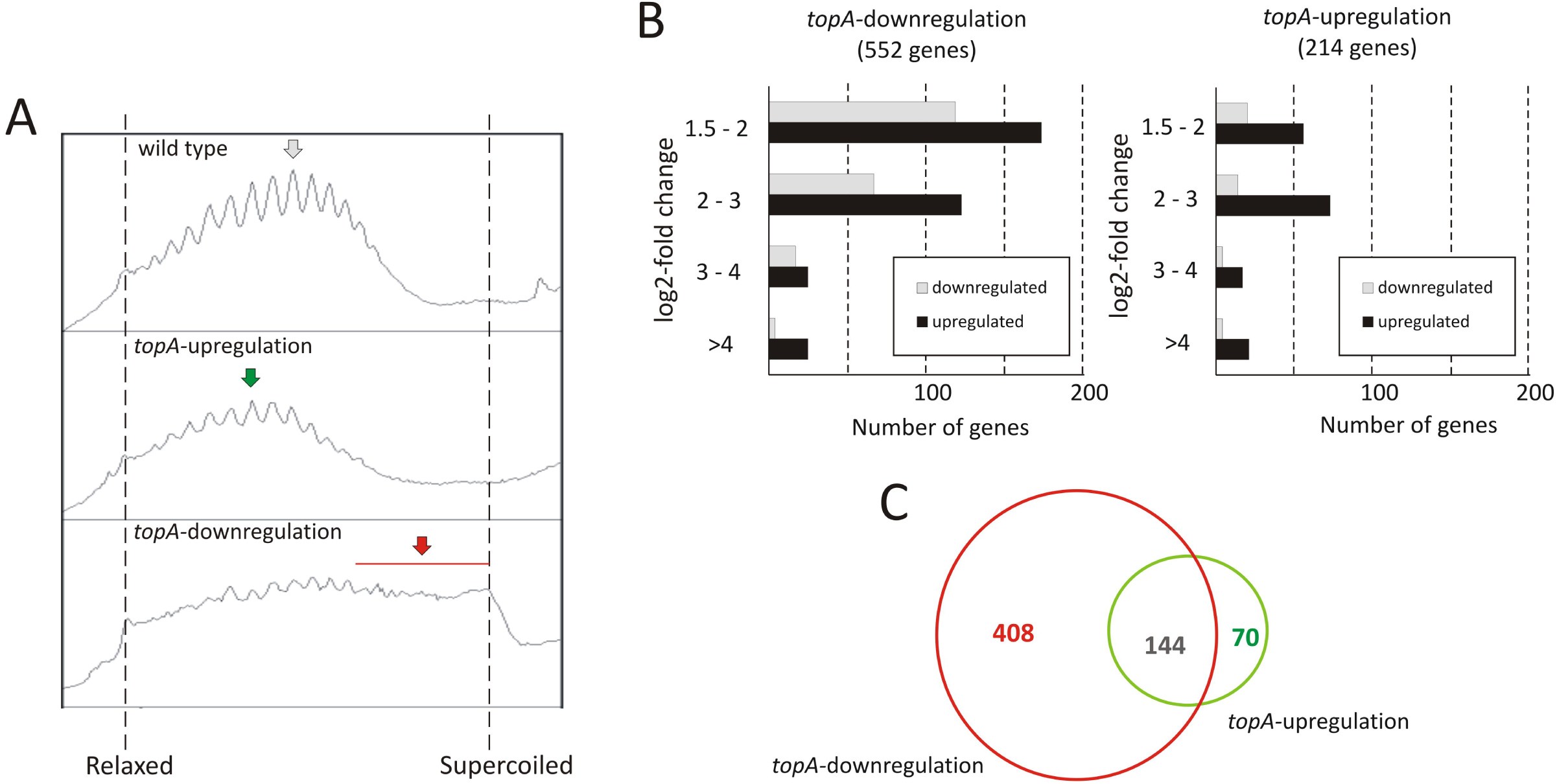
The transcriptional response to long-term supercoiling imbalance. **(A)** The supercoiling density analysis of the reporter plasmid isolated from the wild type strain (upper panel), the *topA*-upregulated strain (middle panel) and the *topA*-downregulated strain (lower panel). The most abundant topoisomers under each condition are marked with arrows. (**B)** Differentially regulated genes identified using RNA-Seq following *topA*-downregulation (left) and *topA*-upregulation (right). The up-(black) or downregulated (gray) genes were grouped according to the magnitude of their transcriptional changes compared to the wild type strain (log2 fold change value). **(C)** Comparison of the number of genes specifically affected by *topA*-upregulation and *topA*-downregulation, and those common to both conditions.

First, we compared the number of *S. coelicolor* genes that were strongly affected by *topA*-down- or upregulation (cultivation in the absence or presence of 1 μg/ml thiostrepton). We identified 552 genes whose transcription was changed by *topA*-downregulation and 214 genes altered by the induction of *topA* transcription (at least 2.83-fold change; log2-fold >1.5), which corresponded to 7.0% and 2.7% of the *S. coelicolor* predicted coding sequences, respectively (Fig. 2B; see supplementary files ‘*topA-downregulation*’ and ‘*topA-upregulation*’ for the complete list of affected genes). Notably, most of the identified ORFs were positively regulated (Fig. 2B), and the positive effect on gene transcription was particularly pronounced following *topA*-upregulation. These observations corroborates with earlier notion that chromosome relaxation preferentially promoted the induction of gene transcription in GC-rich bacteria. Surprisingly, a significant number of genes were sensitive to both changes in supercoiling conditions (Fig. 2C). This may imply that long-term topological imbalances of all types may trigger shared transcriptional response pathway(s). Nevertheless, a substantial fraction of genes responded specifically to increase of negative supercoiling resulting from TopA depletion (408 of 552, more than 70%), whereas *topA*-upregulation regulated specifically only 32% (70 of 214) of identified genes (Fig. 2C). The observed difference in the number of the supercoiling sensitive genes (SSGs), as well as their specificity for particular supercoiling conditions, may be presumably explained by the scale of topological imbalance, which is more pronounced under the former than under the latter conditions (Fig. 2A and S3). However, our results also show that, even though the topological balance was only slightly distorted at *topA* induction, the transcription of a relatively high number of genes was still affected, suggesting the presence of mechanism(s) that maintains a modified level of gene transcription. Markedly, similar observations were made recently for *S. pneumoniae* transcriptional activity of genes under TopA inhibition conditions (Ferrandiz et al., 2016), where the expression of many genes was specifically altered. Collectively, these findings may suggest that transient supercoiling imbalances could lead to long-term rearrangement of bacterial chromosome topology and long-standing changes in gene transcription even after the global supercoiling level is restored.

Overall, we observed that a persistent supercoiling imbalance induced by constitutive change of TopA expression affected the transcription of approximately 3-7% of *S. coelicolor* genes. Interestingly, the inhibition of TopA in *S. pneumoniae* (which was followed by restoring of supercoiling imbalance) affected only 2% of the genome while long term TopA depletion in *S. coelicolor* had more pronounced effect on the gene transcription. This suggests that the response to topological imbalance depends on possibilities to modify transcriptional program, being the most pronounced when the topoisomerase genes expression cannot not be readjusted.

### Long-term supercoiling imbalance impacts the expression of discrete gene clusters

Remarkably, supercoiling sensitive genes whose transcription was affected by the long-term topological imbalance were nonuniformly distributed along the *S. coelicolor* chromosome, similar to genes induced in the novobiocin-treated cells. SSGs induced by *topA*-downregulation or *topA*-upregulation showed characteristic grouping, suggesting they were organized into supercoiling-sensitive clusters, which were in turn separated by supercoiling insensitive regions (Fig. 3A). To quantify these observations, we calculated the percentage of genes affected by different supercoiling conditions within 250 kbp chromosome regions along the whole chromosome. We found that gene clustering was strongly detectable under *topA*-downregulation conditions, with 6-7 chromosomal regions containing more than 5% of genes affected negatively or positively in each cluster (Fig. 3B). Notably, those clusters containing the positively and negatively regulated genes partially overlapped, further supporting the existence of DNA domains that are particularly sensitive to changes in DNA supercoiling. Interestingly, gene clustering was also detectable, although less pronounced, under *topA*-upregulation conditions, where the distribution of the supercoiling-sensitive regions along the chromosome was comparable to that observed in the TopA-depleted strain (Fig. 3B and S4A). Analogous clustering of SSGs has been described previously for *E. coli* and *S. pneumoniae* (Ferrandiz et al., 2016, Peter et al., 2004). Collectively, these observations suggest that the organization of SSGs into topologically separated domains is a feature conserved among bacteria, presumably ensuring the coordinated expression of the associated genes. Remarkably, the loci of the genes affected by topological imbalance showed increased density in the central part of the chromosome, however the initiation of the chromosomal replication region (*oriC*) was found to be outside the identified SSG clusters, and the *oriC* proximal genes, which predominantly encode proteins involved in DNA replication, were insensitive to the changes in chromosome supercoiling (Fig. 3).

**Figure 3.**
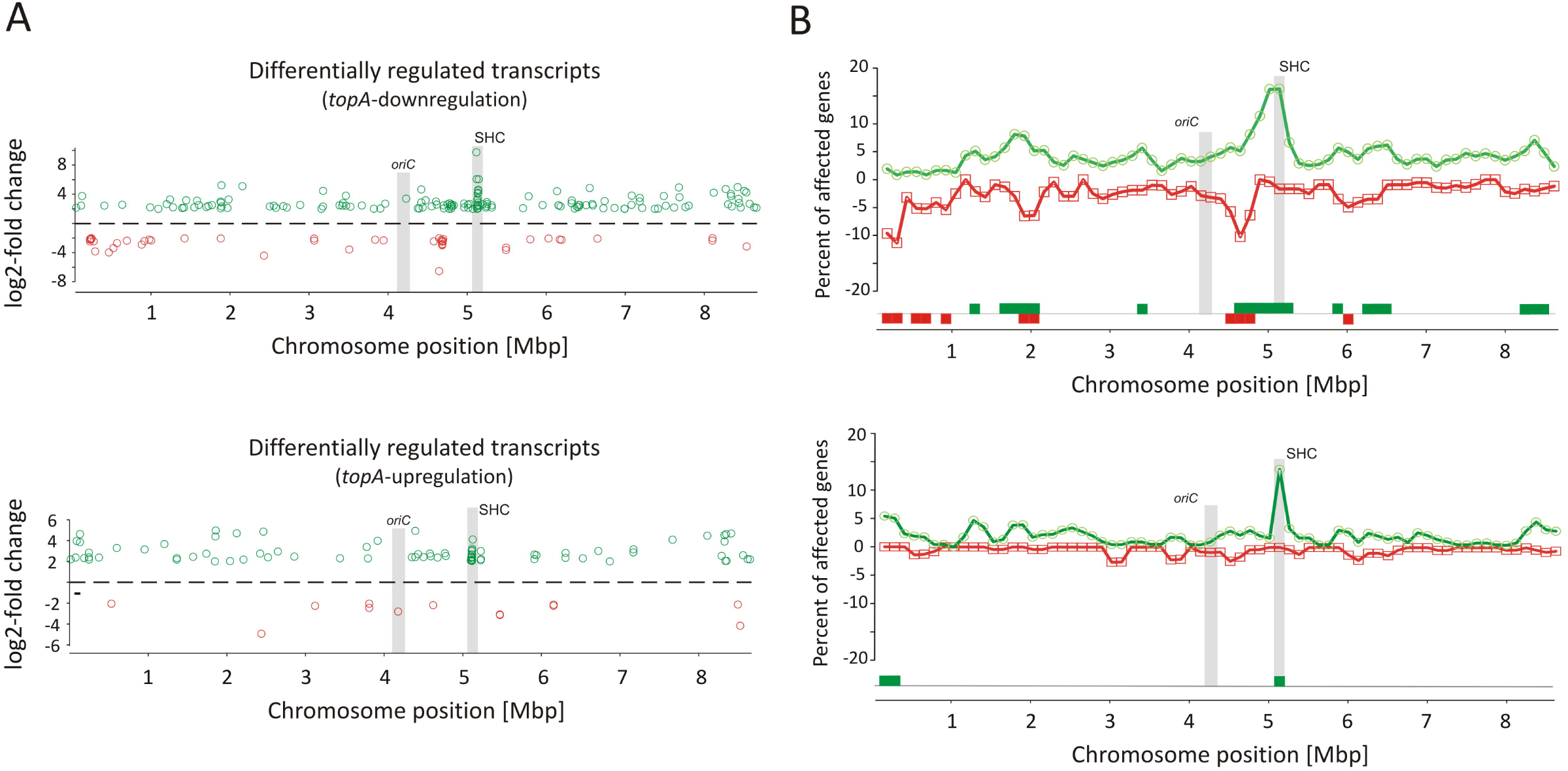
Identification of long-term supercoiling imbalance-sensitive clusters. **(A)** The chromosomal positions of differentially upregulated (green) and downregulated (red) transcripts identified in the *topA*-downregulated strain (upper panel) and *topA*-upregulated strain (lower panel). The positions of the *oriC* region and the supercoiling-hypersensitive cluster (SHC) are marked in gray. **(B)** Identification of supercoiling-sensitive clusters. The percent of genes affected by *topA* downregulation (upper panel) or *topA* upregulation (lower panel) was calculated in a 250-kbp window and the mid of the window was plotted versus chromosome position. The regions with the percentage of upregulated (green) or downregulated (red) genes higher than 5% (supercoiling-sensitive clusters) are marked at the bottom of the plot.

Interestingly, among several identified gene clusters impacted by supercoiling imbalance, one cluster was particularly enriched in SSGs, encompassing ∼20% of all genes positioned within the 250 kbp region. Closer examination of this region revealed that 26 out of 34 genes within 30 kbp (between *sco4667* and *sco4700*) were supercoiling sensitive. We therefore termed this region the <u>s</u>upercoiling-<u>h</u>ypersensitive <u>c</u>luster (SHC). Most of the SHC genes encoded products of unassigned function and these genes themselves were poorly transcribed under standard culture conditions. However, they were strongly upregulated under both *topA*-downregulation and *topA*-upregulation conditions (Table 3). We confirmed these observations and excluded the effect of growth medium or genetic modifications in the PS04 strain, by measuring the relative transcript level in PS04 and control strain cultures grown in liquid 79 medium using RT-qPCR and oligonucleotides specific to the first (*sco4667*) and one before last (*sco4699*) genes of the SHC region. The transcription of both genes appeared to be highly dependent on the level of TopA: their transcription significantly increased when TopA was depleted, decreased when TopA was at the wild type level, and increased slightly again at *topA* induction (Fig. S4B). These observations further supported the proposal that SHC upregulation was correlated with relative degree of topological imbalance. Since we did not observe increased expression of genes in the SHC region following a 10 minute exposure to novobiocin, we speculate that its activation requires not only a supercoiling imbalance, but also the activity of other factors.

**Table 3.** Transcription of genes in the SHC region at constitutive *topA*-down- or upregulation.

| Gene | Locus | Normalized expression* | Log2-fold *topA*-downregulation | Log2-fold *topA*-upregulation | Description |
| --- | --- | --- | --- | --- | --- |
|  | *sco4667* | 12 | 9.75 | 8.93 | two-component system sensor kinase |
|  | *sco4668* | 67 | 7.51 | 6.59 | two-component system response regulator |
|  | *sco4669* | 35 | 6.06 | 5.39 | hypothetical protein |
|  | *sco4670* | 0 | 0.00 | 0.00 | serine protease |
|  | *sco4671* | 21 | 2.31 | 2.08 | LysR family transcriptional regulator |
|  | *sco4672* | 55 | 1.61 | 1.73 | hypothetical protein |
|  | *sco4673* | 56 | 3.20 | 2.35 | DeoR family transcriptional regulator |
|  | *sco4674* | 26 | 2.00 | 2.08 | hypothetical protein |
|  | *sco4675* | 77 | 1.76 | 2.12 | hypothetical protein |
|  | *sco4676* | 3 | 1.22 | 0.41 | hypothetical protein |
| *rsfA* | *sco4677* | 11 | 1.24 | 0.79 | regulatory protein |
|  | *sco4678* | 197 | 1.99 | 1.59 | hypothetical protein |
|  | *sco4679* | 530 | 1.98 | 1.63 | hypothetical protein |
|  | *sco4680* | 116 | 1.37 | 1.56 | DNA-binding protein |
|  | *sco4681* | 137 | 0.69 | 1.29 | short chain dehydrogenase |
|  | *sco4682* | 167 | 0.65 | 1.37 | tautomerase |
| *gdhA* | *sco4683* | 212 | 2.06 | 2.36 | glutamate dehydrogenase |
| *scoF3* | *sco4684* | 3087 | 4.08 | 3.00 | cold shock protein |
|  | *sco4685* | 16 | 3.43 | 2.67 | DEAD/DEAH box helicase |
|  | *sco4686* | 44 | 2.53 | 1.83 | hypothetical protein |
|  | *sco4687* | 674 | 2.38 | 1.79 | hypothetical protein |
|  | *sco4688* | 137 | 2.03 | 1.08 | hypothetical protein |
|  | *sco4689* | 345 | -0.26 | 2.03 | hypothetical protein |
|  | *sco4690* | 148 | 0.41 | 2.39 | hypothetical protein |
|  | *sco4691* | 10 | 1.58 | 2.43 | hypothetical protein |
|  | *sco4692* | 142 | 2.56 | 2.18 | hypothetical protein |
|  | *sco4693* | 62 | 2.86 | 2.03 | hypothetical protein |
|  | *sco4694* | 12 | 4.20 | 2.67 | hypothetical protein |
|  | *sco4695* | 0 | 4.46 | 2.32 | hypothetical protein |
|  | *sco4696* | 8 | 4.62 | 3.22 | hypothetical protein |
|  | *sco4697* | 23 | 4.48 | 3.08 | hypothetical protein |
|  | *sco4698* | 34 | 2.84 | 2.67 | IS1652 transposase |
|  | *sco4699* | 8 | 5.78 | 3.74 | Rhs protein |
|  | *sco4700* | 13 | 6.04 | 4.01 | hypothetical protein |

\* normalization by the upper quartile of gene expression

The SHC genes include those encoding a putative two-component system (*sco4667* and *sco4668*) genes organized in a single operon and additional two putative transcriptional regulator-encoding genes positioned upstream (*sco4671* and *sco4673*). The cluster also encompasses genes whose products may contribute to DNA/RNA transactions, including: putative DNA helicase (*sco4685*), transposase (*sco4698*), and a homolog of *E. coli* Rhs protein (*rearrangement hotspot protein*), which putatively contains the C-terminal toxin domain and is involved in bacterial intercellular competition (Koskiniemi et al., 2013). When we compared the genes in this region to other *Streptomyces* species, we found the synteny was not strong, perhaps suggesting that some of these genes may have been acquired through horizontal transfer (Hindra et al., 2014). This hypothesis is supported by the notion that mobile elements tend to be sensitive to changes in chromosome topology (Lodge and Berg, 1990). Indeed, among 552 affected by TopA depletion conditions we found 16 genes (2.9% of SSGs) encoded putative transposable elements (which constitute only 0.7% of the *S. coelicolor* genome), reinforcing the idea of their increased sensitivity to topological imbalance (see the complete list in the supplementary materials). Why these mobile elements, and more broadly, the genes in the SHC, are specifically subject to topological control, remains an interesting question to address.

In summary, as in other bacteria, in *S. coelicolor* chromosome the supercoiling sensitive domains could be detected. They may constitute the regions of chromosomes particularly prone to changes of supercoiling and/or may be regulated by supercoiling sensitive transcriptional regulators.

### Long-term topological imbalance affects the transcription of genes encoding topology-controlling proteins

Since topological homeostasis could not be restored in the TopA-depleted *S. coelicolor* PS04 strain, either by induction of the *topA* promoter or by *gyrBA* silencing (Szafran et al., 2016), we predicted that the resulting increase in DNA supercoiling may promote changes in the level of NAPs to compensate for the changes in chromosome topology. According to previous reports (Bradshaw et al., 2013) NAPs are present at high levels during wild type *S. coelicolor* vegetative growth. We confirmed that under standard growth conditions, many NAP genes, including *hupA*, *ssbA*, *lsr2*, *sihf* and *hupS* were highly transcribed (Fig. 4A), while other sporulation and/or stress responsive genes like *dpsB*, *dpsC* and *ssbB*, were poorly transcribed. Next we assessed the impact of the chromosome supercoiling changes on the expression of these genes.

**Figure 4.**
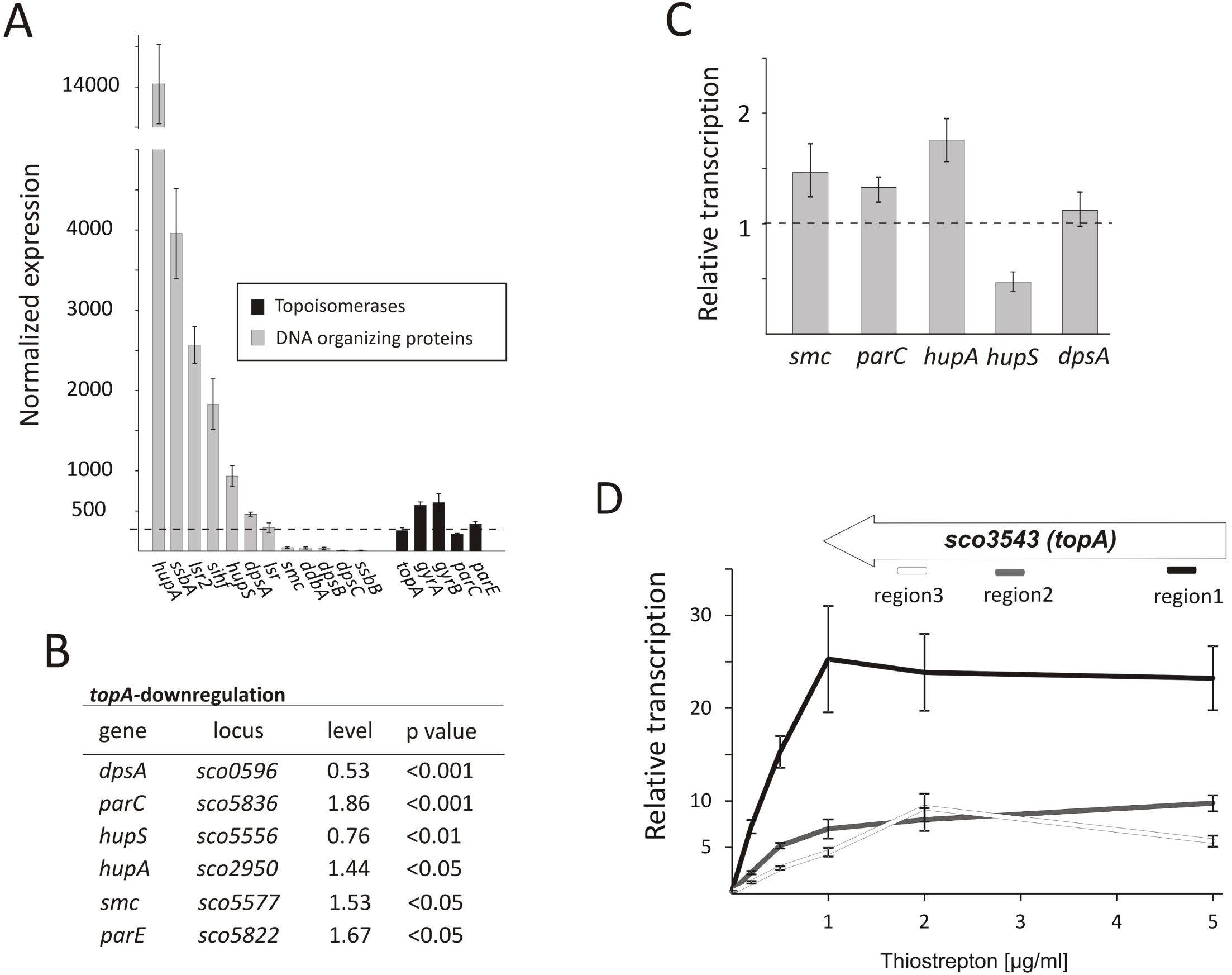
The topological imbalance-induced transcriptional response of the genes encoding chromosome organizing proteins. **(A)** RNA-Seq based expression of the genes encoding the DNA-organizing proteins (gray) and topoisomerases (black) in the wild type *S. coelicolor* strain normalized by the upper quartile. The error bars correspond to standard deviations calculated for 4 independent biological replicates. **(B)** RNA-Seq analysis of the relative transcriptional response of genes encoding DNA-organizing proteins under *topA*-downregulation conditions. The expression level of particular genes in the wild type strain was estimated as 1. **(C)** The relative transcription of selected genes in the TopA-depleted and control wild type strains calculated using RT-qPCR analysis. **(D)** RT-qPCR comparison of nonuniform *topA* gene transcription under different concentrations of *topA* inducer using oligonucleotide pairs specific for particular transcript regions. The transcript enrichment of particular regions was normalized by the expression of the *hrdB* gene which served as the endogenous control and compared to the wild type strain, in which transcription was estimated as 1. The amplified regions of the *topA* gene (1- white; 2 - gray, and 3 - black) are shown above.

Following TopA-depletion (conditions of increased negative supercoiling), we observed transcript levels of DNA-organizing proteins to be statistically different (p < 0.05) (Fig. 4B). Of the NAPs affected by changes in chromosome topology, *hupA* and *hupS* were amongst the most impacted. Notably, however, the transcriptional effects were in opposite directions: *hupA* transcript levels increased to 1.44 relative to wild type conditions, while *hupS* transcript levels decreased to 0.76 compared with wild type (Fig. 4B). To assess whether these effects were growth medium-specific, and to confirm the supercoiling dependence of these two HU-encoding genes, we grew these strains in the rich 79 medium, and using RT-qPCR compared their expression under a range of *topA* induction conditions. As before, we observed *hupA* upregulation and *hupS* downregulation following TopA depletion if compared to the wild type level (Fig. 4C). Previous work has suggested that HupS functions primarily during *S. coelicolor* sporulation (Salerno et al., 2009); however, both HU homologs, HupS and HupA, were shown to be present during *S. coelicolor* vegetative growth, although their levels are unequal (Bradshaw et al., 2013) (our transcriptional analysis showed that *hupA* transcript level is over 14 times higher than *hupS* transcript). Our observations suggest that both *S. coelicolor* HU homologs may contribute to the interplay of different factors that facilitate adjustment to altered supercoiling conditions. In *E. coli* two HU homologs, HupA and HupB form homo- and heterodimers at different growth stages, with the heterodimer having a crucial role when the culture reaches stationary phase (Claret and Rouviere-Yaniv, 1997). Furthermore, DNA supercoiling influences *hupB* transcription in *Mycobacteria*, and its product affects TopA activity (Ghosh et al., 2014; Guha et al., 2018), suggesting that cooperation interactions between HU proteins and TopA may be conserved in bacteria.

In addition to the effects on *hupA* and *hupS*, increased chromosome supercoiling also profoundly affected the transcription of *dpsA* (decreased to 0.53 of wild type levels), *smc* (increased to 1.53 of wild type levels) (Fig. 4B) and two genes (*parC* and *parE*) encoding topoisomerase IV subunits (increased to 1.86 and 1.67 of wild type levels, respectively). Using RT-qPCR, we confirmed upregulation of *parC*, *parE* and *smc* genes. Similar observations have been made in *E. coli*, where increases in negative supercoiling result in the induction of *mukBEF* expression (Peter et al., 2004), where these genes encode homologs of SMC. Additionally, it is conceivable, that upregulating the expression of *parC* and *parE* (topoisomerase IV) may partially complement the effects of TopA depletion. However, the changes in *dpsA* transcription were not confirmed using RT-qPCR in cultures grown in different medium than used in RNA-Seq experiment (Fig. 4B and 4C). A possible explanation for this observation is that *dpsA* transcription is affected by DNA supercoiling but only under osmotic stress conditions (in YEME/TSB medium used in RNA-Seq experiment but not in the rich 79 medium used for RT-qPCR experiment)(Facey et al., 2009). Whether and how the changes in expression of genes encoding DNA organizing proteins influence chromosome topology have yet to be determined.

Next, we assessed the effect of elevated *topA* expression (15-25x higher transcript levels compared with levels in the wild type strain). As mentioned, these increased *topA* transcript level was not correlated with increased TopA protein which level was elevated only 22% in comparison with the wild type strain ((Szafran et al., 2016) Fig. 1 and S3A), thus only slightly affecting DNA supercoiling. Based on that observation, we speculated that the TopA may be regulated posttranscriptionally and/or posttranslationally (e.g. may be subject to transcript processing or specific protein degradation). Indeed, while *topA* transcript levels were uniformly distributed along the 2858 bp *topA* gene in the wild type and the *topA*-downregulation strain), *topA* induction considerably increased the fraction of reads mapping to the beginning (region1) but not to middle (region2) or end (region3) of the *topA* gene (Fig. S5A). The nonuniform *topA* transcript level was confirmed using RT-qPCR with the specific oligonucleotides complementary to the distinct fragments of the *topA* gene (Fig. 4D). Interestingly, in the closely related *M. smegmatis*, gyrase binds within the *topA* promoter region, preventing its transcription (Ahmed et al., 2015) thus, similar mechanism of *topA* autoregulation in *Streptomyces* cannot be excluded as well, however, at the moment it remains only speculative. On the other hand, the possibility that *topA* transcripts are also subject to posttranstriptional processing and/or mRNA degradation can also explain the nonuniform detection of the particular *topA* transcript regions.

In summary, the increased level of negative DNA supercoiling induces the coordinated modification of topoisomerases and NAPs level in attempt to restore the topological balance. Among them, the HupA protein may play a crucial role in maintaining nucleoid architecture under TopA depletion; with potential contributions from topoisomerase IV (ParCE) and downregulation of HupS – the other HU homolog present in *Streptomyces*. Intriguingly, *topA* induction had little effect on global DNA supercoiling, as the *Streptomyces* appear to have mechanisms in place to ensure TopA levels are maintained below a certain threshold, through possible autoregulatory control.

### Alternating *topA* transcript levels stimulates the expression of DNA damage response genes

Chromosome regulation stemming from gyrase inhibition with novobiocin had an immediate impact on 1.5% of the genes in the *S. coelicolor* chromosome. Interestingly, among the genes affected by severe depletion or modest increase of TopA level, we identified genes whose transcription was modified under both conditions.

Among the genes sensitive to any alteration of TopA level, 14 regulatory protein-encoding genes were identified (Table 4), including the four regulators in the SHC region (*sco4667* (sensor kinase), *sco4668* (regulatory protein), and putative transcriptional regulators (*sco4671* and *sco4673*)). Outside of this region was *sco5803*, which encodes the LexA repressor that controls the DNA damage response (highly induced by *topA*-up- and downregulation). In *E. coli*, the LexA-regulon encompasses at least 31 genes, including *recA* and *lexA* (the latter being negatively autoregulated)*, uvrA, ftsK, polB, dinF*, and *dnaE2*, alongside others involved in the DNA damage response (Fernandez De Henestrosa et al., 2000). We found that the transcription of *recA*, *recX*, *dnaE2*, *dinP* or *uvrA* genes, which presumably belong to the LexA-regulon in *S. coelicolor*, based on the predicted binding consensus sequence (Novichkov et al., 2013), were similarly affected by changes in chromosome supercoiling, indicating that the entire LexA regulon was impacted (Table 5 and 6 and Fig. S6).

**Table 4.** The regulators encoding genes affected by both constitutive *topA*-downregulation and *topA*-upregulation.

| Gene | Locus | Normalized expression* | Log2-fold *topA*-downregulation | Log2-fold *topA*-upregulation | Description |
| --- | --- | --- | --- | --- | --- |
| **REGULATORS** |  |  |  |  |  |
|  | *sco3435* | 14 | 1.72 | 1.58 | transcriptional regulator |
|  | *sco4214* | 2429 | -6.55 | -6.60 | AbaA-like regulatory protein |
| *xlnR* | *sco4215* | 820 | -2.05 | -2.20 | GntR family transcriptional regulator |
|  | *sco4671* | 21 | 2.31 | 2.08 | LysR family transcriptional regulator |
|  | *sco4673* | 56 | 3.20 | 2.35 | DeoR family transcriptional regulator |
|  | *sco5656* | 2804 | -1.80 | -2.15 | transcriptional regulator |
| *lexA* | *sco5803* | 240 | 2.04 | 1.62 | LexA repressor |
| *nsdB* | *sco7252* | 118 | -1.67 | -2.80 | regulatory protein |
| **TCSs** |  |  |  |  |  |
|  | *sco4667* | 12 | 9.75 | 8.93 | two-component system sensor kinase |
|  | *sco4668* | 67 | 7.51 | 6.59 | two-component system response regulator |
| **PROTEASES** |  |  |  |  |  |
| *Lon* | *sco5285* | 10 | 1.89 | 1.85 | ATP-dependent protease |
| **GNATs** |  |  |  |  |  |
|  | *sco1624* | 13 | 2.51 | 2.74 | acetyltransferase |
|  | *sco2379* | 12 | 2.14 | 1.87 | acetyltransferase |
| *desC* | *sco2784* | 285 | -2.32 | -1.83 | acetyltransferase |

\* normalization by the upper quartile of gene expression

**Table 5.** DNA processing and DNA repair genes impacted by constitutive *topA*-downregulation.

| Gene | Locus | Normalized expression* | Log2-fold change | Description |
| --- | --- | --- | --- | --- |
| *dnaE2* | *sco1739* | 5 | 5.22 | DNA polymerase III subunit alpha |
| *dinP* | *sco1738* | 6 | 4.84 | DNA polymerase IV |
|  | *sco1767* | 29 | 1.92 | DNA hydrolase |
|  | *sco1827* | 72 | 1.51 | DNA polymerase III subunit epsilon |
| *uvrA* | *sco1958* | 69 | 1.89 | exonuclease ABC subunit A |
|  | *sco1969* | 67 | 1.88 | DNA-methyltransferase |
|  | *sco2863* | 18 | 1.98 | helicase |
|  | *sco3434* | 27 | 2.01 | DNA polymerase I |
|  | *sco4685* | 16 | 3.43 | DEAD/DEAH box helicase |
| *xseA* | *sco5056* | 69 | 1.51 | exodeoxyribonuclease VII large subunit |
|  | *sco5183* | 12 | 1.70 | ATP-dependent DNA helicase |
|  | *sco5184* | 11 | 1.54 | ATP-dependent DNA helicase |
| *recX* | *sco5770* | 136 | 3.15 | recombination regulator RecX |
| *recA* | *sco5769* | 426 | 2.76 | recombinase A |
| *lexA* | *sco5803* | 240 | 2.04 | LexA repressor |
|  | *sco5920* | 9 | 1.53 | DEAD/DEAH box helicase |
|  | *sco6719* | 2 | 3.86 | UvrA-like ABC transporter |
|  | *sco7522* | 7 | 2.32 | DNA ligase |

\* normalization by the upper quartile of gene expression

**Table 6.** DNA processing and DNA repair genes impacted by constitutive *topA*-upregulation.

| Gene | Locus | Normalized expression* | Log2-fold change | Description |
| --- | --- | --- | --- | --- |
| *dinP* | *sco1738* | 6 | 5.04 | DNA polymerase IV |
| *dnaE2* | *sco1739* | 5 | 4.96 | DNA polymerase III subunit alpha |
|  | *sco1767* | 122 | 1.81 | DNA hydrolase |
| *uvrA* | *sco1958* | 69 | 1.98 | excinuclease ABC subunit A |
|  | *sco3434* | 27 | 2.29 | DNA polymerase I |
|  | *sco3798* | 21 | -2.81 | chromosome condensation protein |
|  | *sco4685* | 16 | 2.67 | DEAD/DEAH box helicase |
|  | *sco5761* | 20 | 1.54 | ATP-dependent DNA helicase |
| *recA* | *sco5769* | 426 | 2.34 | Recombinase A |
| *recX* | *sco5770* | 136 | 2.81 | recombination regulator RecX |
| *lexA* | *sco5803* | 240 | 1.62 | LexA repressor |
|  | *sco7522* | 7 | 2.05 | DNA ligase |

\* normalization by the upper quartile of gene expression

The identification of putative LexA-dependent genes among the genes upregulated when *topA* transcription was altered may suggest that these conditions trigger LexA-dependent DNA repair pathway. However, the mechanism of LexA regulon induction by altered chromosome supercoiling remains speculative. We assume that in *Streptomyces*, as in other bacteria, LexA activity is controlled by RecA, which in the presence of ssDNA and in an ATP-dependent manner stimulates the self-cleavage of LexA, resulting in derepression of the LexA regulon. It is conceivable that changes in chromosome topology alter LexA binding to DNA, or that the formation of single-stranded DNA patches upon depleting TopA (Parsa et al., 2012) promote RecA-dependent cleavage of LexA and derepression of LexA-regulated genes. However, the latter possibility could not account for the release of LexA repression during chromosome relaxation and further studies are required to understand the mechanism underlying LexA regulation in response to supercoiling changes. Since DNA damage is frequently accompanied by changes in local chromosome architecture, it is possible that changes in chromosome supercoiling may be sensed as a sign of DNA damage, leading to the induction of DNA repair genes. Interestingly, in *Corynebacterium glutamicum*, an actinobacterial relative of *Streptomyces*, induction of the DNA repair pathway was associated with the inhibition of cell division, which could explain the inhibition of sporulation seen for the *S. coelicolor* TopA-depleted strain (Ogino et al., 2008).

Since we did not identify LexA-regulon among novobiocin-sensitive genes, we speculate that the DNA repair response may be the effect of long-term exposure to topological imbalance. Thus, topological imbalance if cannot be compensated by response pathways could serve as a marker of DNA damage.

### Many transcriptional regulators are sensitive to specific supercoiling conditions

Although *topA* downregulation and *topA*-upregulation induced a common transcriptional response, a significant fraction of the affected genes responded specifically to particular supercoiling conditions (Fig. 2C). We identified 12 SSGs-encoding regulatory proteins that were affected specifically by *topA*-upregulation (Table 7). These numbers are similar to those induced by novobiocin treatment, but surprisingly, the sets of genes identified in both experiments did not overlap. Amongst the many uncharacterized genes that were sensitive to *topA*-upregulation, developmental regulators such as *sigF* (upregulated) and *nsdA* (downregulated) were identified; however, these changes in expression were not associated with any obvious phenotype for the *topA* up-regulated strain.

**Table 7.** Regulatory protein-encoding genes impacted by constitutive *topA*-upregulation.

| Gene | Locus | Normalized expression* | Log2-fold change | Description |
| --- | --- | --- | --- | --- |
| **REGULATORS** |  |  |  |  |
|  | *sco0140* | 3 | 3.84 | MerR family transcriptional regulator |
|  | *sco2865* | 154 | -2.27 | regulatory protein |
| *sigF* | *sco4035* | 4 | 2.70 | RNA polymerase sigma factor |
|  | *sco4102* | 6 | 2.42 | MerR family transcriptional regulator |
|  | *sco5025* | 22 | 7.71 | transcriptional regulator |
|  | *sco5437* | 41 | 2.65 | MerR family transcriptional regulator |
| *nsdA* | *sco5582* | 67 | -1.67 | Regulator |
|  | *sco5962* | 2 | 8.10 | transcriptional regulator |
|  | *sco6520* | 34 | -1.63 | RNA polymerase sigma factor |
|  | *sco7530* | 6 | 4.61 | regulatory protein |
|  | *sco7698* | 2 | 9.13 | MerR family transcriptional regulator |
| **TCSs** |  |  |  |  |
| *cutS* | *sco5863* | 116 | -1.57 | two-component sensor kinase |

\* normalization by the upper quartile of gene expression

The increased chromosome supercoiling resulting from TopA depletion specifically influenced a substantial number of genes encoding regulatory proteins. Among 35 of the TopA depletion-sensitive genes, we identified 18 putative transcriptional regulators, including seven genes encoding TCSs (kinases and/or their putative phosphorylation targets), six genes encoding putative acetyltransferases (GNATs family) and four genes encoding subunits of regulatory Clp proteases (Table 8), alongside the sporulation-specific sigma factor encoding *whiG*.

**Table 8.** Regulatory genes affected by *topA*-downregulation.

| Gene | Locus | Normalized expression* | Log2-fold change | Description |
| --- | --- | --- | --- | --- |
| **REGULATORS** |  |  |  |  |
|  | *sco0132* | 56 | -2.49 | transcriptional regulator |
|  | *sco0204* | 1059 | -1.88 | LuxR family transcriptional regulator |
|  | *sco1104* | 56 | 1.74 | TetR family transcriptional regulator |
|  | *sco1736* | 6 | 2.84 | MarR family transcriptional regulator |
|  | *sco2223* | 54 | 1.72 | TetR family transcriptional regulator |
|  | *sco4059* | 22 | 1.61 | transcriptional regulator |
|  | *sco4375* | 9 | 1.58 | MarR family regulatory protein |
|  | *sco5323* | 52 | 1.57 | transcriptional regulator AsnC |
|  | *sco5418* | 8 | 1.81 | transcriptional regulator |
| *whiG* | *sco5621* | 6 | 4.40 | RNA polymerase sigma factor WhiG |
|  | *sco5840* | 11 | 2.24 | transcriptional regulator |
|  | *sco6299* | 35 | 1.57 | TetR family transcriptional regulator |
|  | *sco6664* | 14 | 2.03 | transcriptional regulator |
|  | *sco7014* | 22 | 1.75 | LacI family transcriptional regulator |
|  | *sco7042* | 114 | 1.60 | MarR family transcriptional regulator |
|  | *sco7270* | 15 | 1.62 | transcriptional regulator |
|  | *sco7639* | 45 | 1.84 | MarR family regulatory protein |
|  | *sco7727* | 100 | -1.56 | MarR family regulatory protein |
| **TCSs** |  |  |  |  |
| *ohkA* | *sco1596* | 31 | 1.77 | two-component sensor kinase |
| *abrA2* | *sco1745* | 7 | 1.89 | two-component system response regulator |
|  | *sco3144* | 6 | 2.42 | two-component system response regulator |
|  | *sco3818* | 179 | -1.96 | two-component system response transcriptional regulator |
|  | *sco4362* | 8 | 2.29 | two component system sensor kinase |
|  | *sco5455* | 15 | 1.87 | two-component system response regulator |
|  | *sco7649* | 15 | 1.65 | two-component system sensor kinase |
| **PROTEASES** |  |  |  |  |
| *clpP2* | *sco2618* | 2584 | -1.77 | ATP-dependent Clp protease proteolytic subunit |
| *clpP1* | *sco2619* | 3091 | -1.61 | ATP-dependent Clp protease proteolytic subunit |
| *clpA* | *sco6408* | 29 | -1.69 | Clp protease ATP-binding subunit |
| *clpP4* | *sco7280* | 8 | 1.91 | ATP-dependent Clp protease proteolytic subunit 2 |
| **GNATs** |  |  |  |  |
|  | *sco1563* | 46 | 1.55 | Acetyltransferase |
|  | *sco2227* | 33 | -2.24 | Acetyltransferase |
|  | *sco2876* | 14 | 2.00 | Acetyltransferase |
|  | *sco4050* | 93 | -2.15 | Acetyltransferase |
|  | *sco5228* | 8 | 2.09 | Acetyltransferase |
|  | *sco7447* | 1 | 4.70 | Acetyltransferase |

\* normalization by the upper quartile of gene expression

Transcription of the gene encoding the sigma factor WhiG was one of the most significantly upregulated regulatory genes under TopA depletion (over 21-fold induction; Table 8 and Fig. 5A). RT-qPCR confirmed that *whiG* transcription was stimulated by increased DNA supercoiling and decreased to wild type levels upon induction of normal *topA* transcription (Fig. 5B). WhiG is typically expressed at low levels during vegetative growth and is involved in the regulatory cascade that governs *Streptomyces* sporulation (Elliot et al., 2001; Kelemen et al., 1996). WhiG directs the expression of the *whiI* and *whiH* genes, but does not affect the transcription of other *whi*-family regulators, such as *whiA* and *whiB* (Chater et al., 1989; Mendez and Chater, 1987; Ryding et al., 1998), we analyzed the transcriptional response of these genes to increased DNA supercoiling. As expected, the transcription of both *whiI* and *whiH* was elevated following TopA depletion (although they were not identified in the initial screening due to below-threshold q values, remarkably p <0.001 was still statistically relevant), suggesting the induction of a WhiG-dependent regulatory cascade by changes in chromosome supercoiling (Fig. 5A). In *S. coelicolor*, in contrast to *S. venezuelae*, *whiG* transcription was independent of any of six known *whi* genes (*whiA*, *B, G, H, I* and *J*) (Kelemen et al., 1996). However, *whiG* was shown to belong to the BldD-regulon, which also encompasses *bldN*, *bldM*, and *whiB* (den Hengst et al., 2010; Elliot et al., 2001). In fact, in *S. venezuelae*, the *whiG* gene was shown to be directly repressed by BldD, the master regulator that binds to DNA upon association with c-di-GMP (Tschowri et al., 2014). This prompted us to test whether increased DNA supercoiling influenced the transcription of other BldD-dependent genes. Indeed, upon TopA depletion, we observed high upregulation of three BldD targets: *bldM*, *bldN* and *whiG* (Fig. 5A). The mechanism of supercoiling-dependent upregulation of the BldD-regulon may, similar to that proposed for LexA, result from decreased BldD DNA-binding affinity during higher DNA supercoiling. Thus, alleviating BldD binding during times of increased DNA supercoiling could lead to increased target gene transcription.

**Figure 5.**
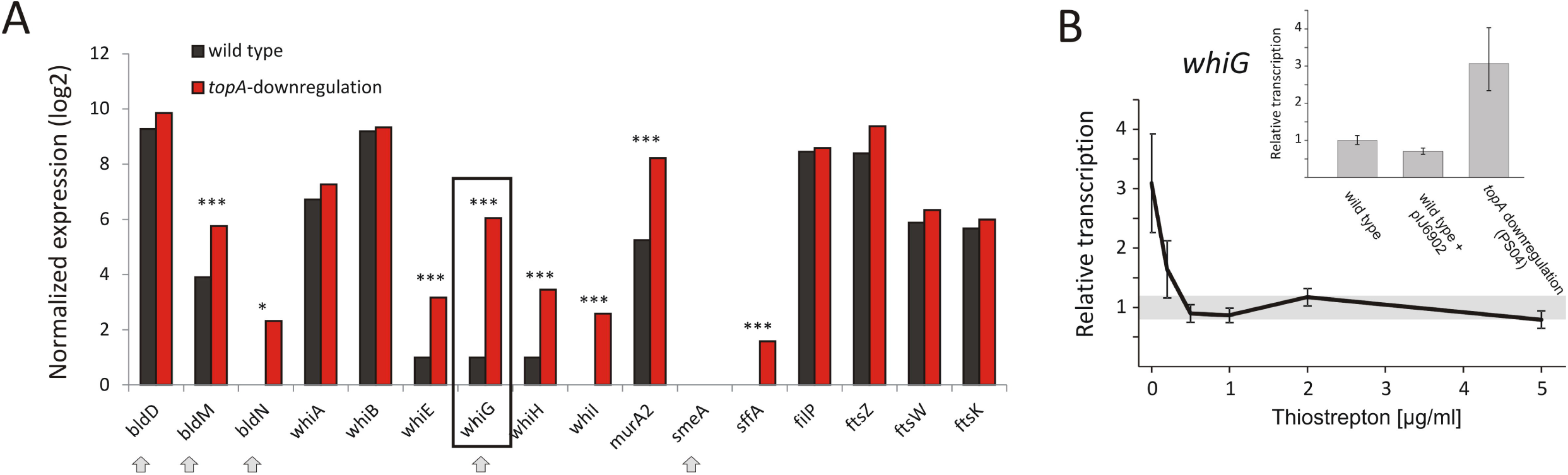
TopA depletion-induced changes in the transcription of genes involved in sporulation regulatory cascades. **(A)** RNA-Seq based expression of selected genes encoding for *whi* and *bld* regulatory cascades genes normalized by the upper quartile in the wild type strain (black) and the TopA-depleted strain (red). The genes belonging to the BldD regulon are marked with arrows. The statistically significant changes in transcript levels were marked with asterisks (* p value < 0.05; *** p value <0.001). p values were calculated based on RNA-Seq data analysis for 4 independent biological replicates for the wild type and *topA*-downregulated strains. **(B)**e RT-qPCR analysis of *whiG* transcript levels at different concentrations of the *topA* gene inducer. wild type levels of *whiG* transcription were set at 1, and standard deviation for wild type transcription is marked with a gray bar. The inset shows RT-qPCR analysis of *whiG* transcript levels in the TopA-depleted and the wild type strains, as well as the control strain that contained the pIJ6902 plasmid without the *topA* gene. The *whiG* transcript enrichment was normalized by the expression of the *hrdB* gene which served as the endogenous control and compared to the wild type strain in which transcription was estimated as 1.

Remarkably, even though elevated supercoiling induced key sporulation genes including *whiG*, their induction did not lead to sporulation; in fact the TopA-depleted strain fails to form spores (Szafran et al., 2013). This may indicate either a lack of additional regulators needed for differentiation, or the activity of other signaling pathways that prevent sporulation (*e.g*. the DNA damage response, as described for RecA/LexA above). The observed induction of sporulation cascades does, however, suggest that DNA topology may function as a global regulator that triggers sporulation cascades either in response to environmental stress or physiological conditions that induce increased DNA supercoiling.

## SUMMARY

Topological imbalance has profound effects on gene transcription in all bacteria; however, different organisms evoke distinct responses (Ferrandiz et al., 2010; Gmuender et al., 2001; Peter et al., 2004). We showed here that in *S. coelicolor* supercoiling-sensitive genes are organized in discrete clusters, a feature that seems to be a conserved strategy for topological regulation among many bacteria. Supercoiling imbalance triggers the activation of genes involved in stress responses, including DNA repair pathway, transmembrane transporters and chaperonins, as well as genes involved in the production of secondary metabolites. Both increased and decreased DNA supercoiling appear to have been detected as signals of DNA damage, inducing DNA repair genes and oxidative stress response genes. The long-term response to supercoiling imbalance is based on the activation of the set of primary and downstream regulatory proteins. Sporulation is triggered under stress conditions in *Streptomyces*, and accordingly, we found that several differentiation-regulating genes are affected by DNA supercoiling. In general, topology-governed regulons are expected to rely on the supercoiling-dependent binding of various regulatory proteins to DNA.

## Supporting information

novobiocin

topA-downregulation

topA-upregulation

## FUNDING

This study was supported by: The National Science Center, Poland: OPUS grant 2014/15/NZ2/01067 to DJ; HARMONIA grant 2016/22/NZ1/00122 to MJS and PRELUDIUM grant 2016//23/N/NZ2/01169 to MG

The Canadian Institute of Health Research (CIHR), and Natural Sciences and Engineering Research Council (NSERC) to ME

