## Supplementary Figures for "Transcriptional response of *Streptomyces coelicolor* to rapid chromosome relaxation or long-term supercoiling imbalance"

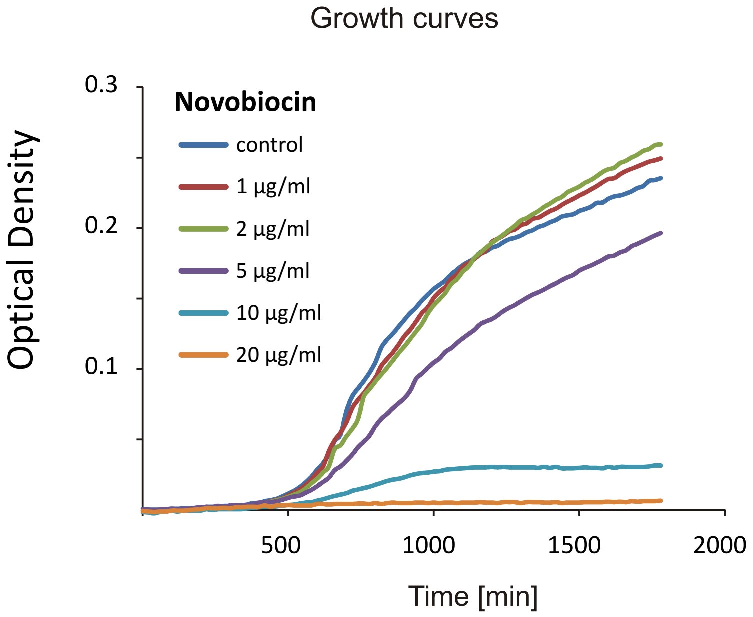


**Figure S1. *S. coelicolor* growth in presence of novobiocin.** The wild type *S. coelicolor* (M145) was cultured in the presence of increasing concentrations of novobiocin (0 - 20 µg/mL) in liquid 79 medium at 30°C. The culture growth was monitored using the Bioscreen C instrument. The optical density of the culture was measured at 600 nm wavelength in 20-minute intervals. Each conditions were analyzed in triplicate and subsequently averaged values were plotted against time of culture.

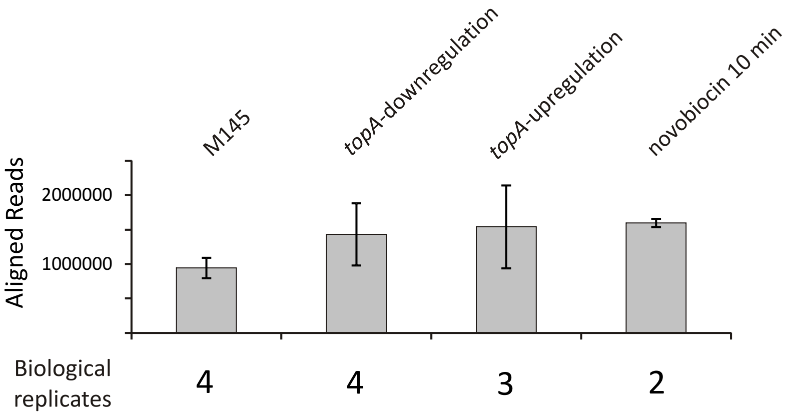


**Figure S2. The number of aligned reads in the RNA-Seq experiments.** The reads obtained from MiSeq Illumina sequencing of the libraries for the wild type strain as well as the strains with modified DNA supercoiling conditions (*topA* gene upregulation and downregulation, as well as the wild type strain exposed for 10 minutes to novobiocin 10 µg/ml) were aligned using Rockhopper software. Below the diagram, the number of

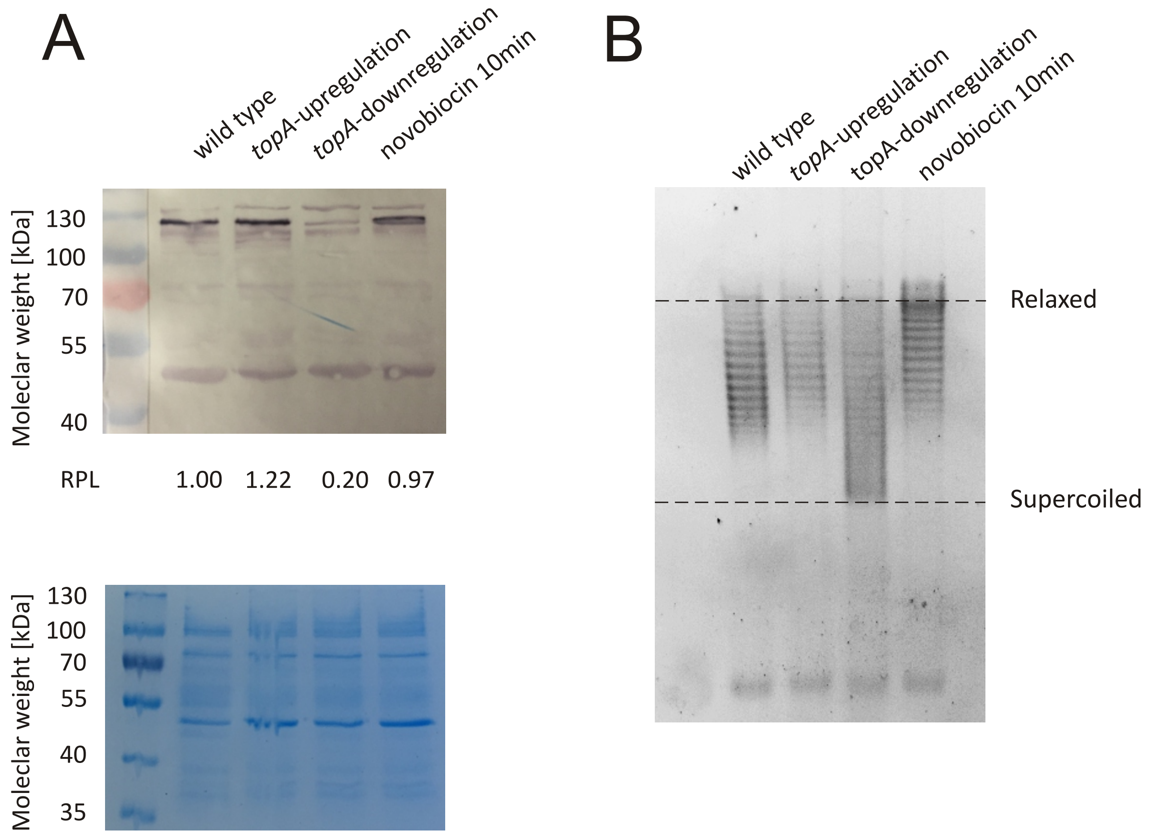


**Figure S3. TopA levels and DNA supercoiling under different experimental conditions.** (**A**) Western blot analysis of *S. coelicolor* TopA levels using anti-TopA polyclonal antibodies (upper panel). The loading control for the experiment (lower panel). The relative protein level (RPL) was calculated as the TopA band intensity in comparison to the wild type TopA level estimated as 1.00 (**B**) Plasmid supercoiling assay, in which the reporter plasmid was isolated from the control strain (MS10) and the strain with modified *topA*-transcription (MS11) (upregulation and downregulation of the *topA* gene) or the control strain (MS10) treated with novobiocin.

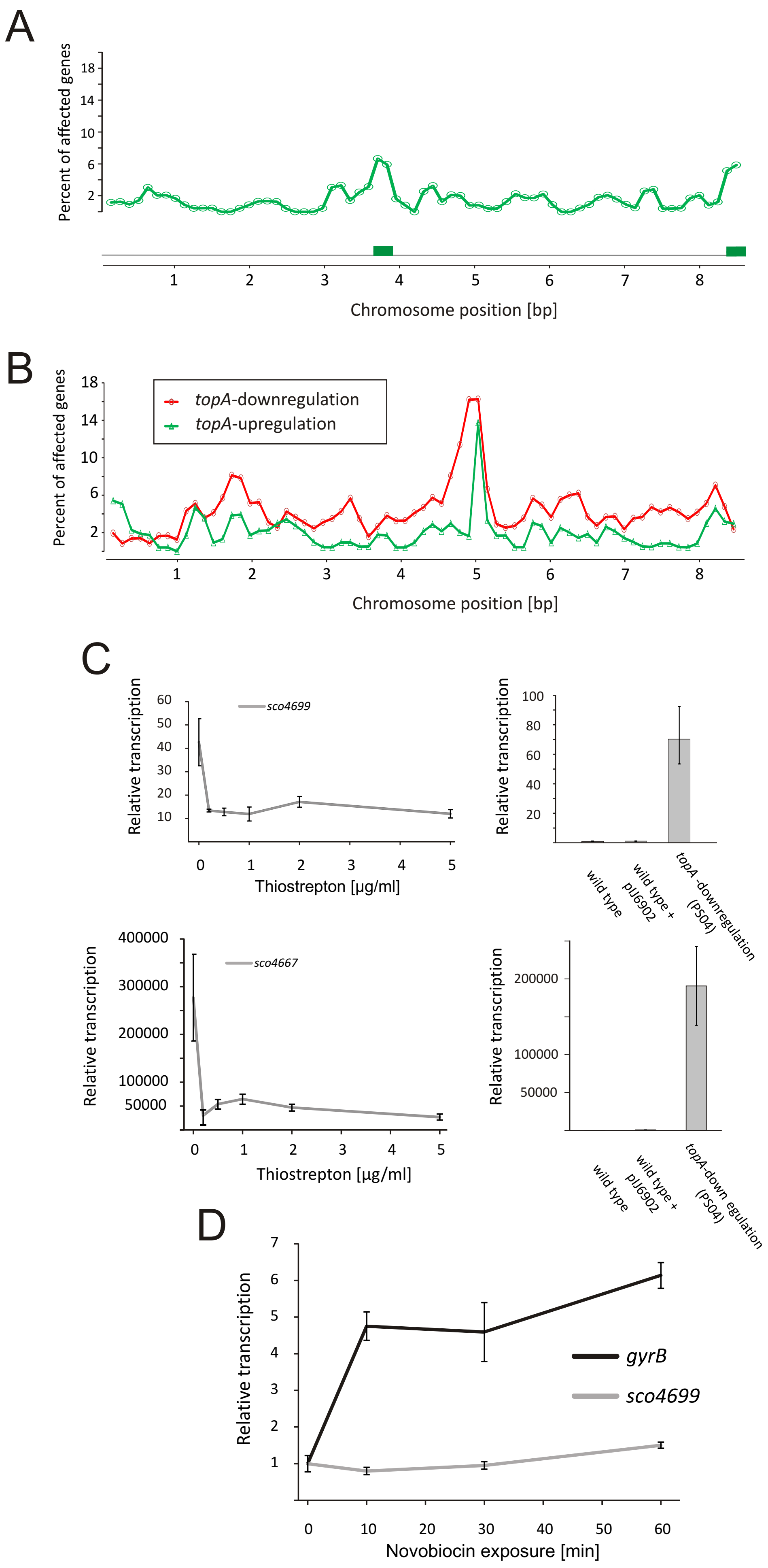
A
Percent of affected genes
18
14
10
6
2
1
2
3
4
5
6
7
8
Chromosome position [bp]
B
Percent of affected genes
topA-downregulation
topA-upregulation
Chromosome position [bp]
C
Relative transcription
sco4699
60
50
40
30
20
10
0
1
2
3
4
5
Thiostrepton [µg/ml]
100
80
60
40
20
wild type
wild type + pIJ6902
topA -downregulation (PS04)
sco4667
400000
300000
200000
100000
50000
wild type
wild type + pIJ6902
topA-down egulation (PS04)
D
7
6
5
4
3
2
1
gyrB
sco4699
0
10
20
30
40
50
60
Novobiocin exposure [min]

**Figure S4. Supercoiling-sensitive clusters.** (**A**) Identification of novobiocin-sensitive clusters. The percent of genes upregulated in response to novobiocin was calculated in 250-kbp windows (with a sliding window of 125 kbp) and subsequently the mid of the window was plotted versus chromosome position. The regions containing more than 5% of the upregulated genes are indicated in green and are deemed to be novobiocin-sensitive clusters. (**B**) Overlay of the plots showing the percentage of upregulated genes in the 250-kbp genome fragment identified in the *topA*-downregulated and *topA*-upregulated strains. (**C**) RT-qPCR analysis of *sco4699* (upper panel) and *sco4667* (lower panel) transcription at different concentrations of *topA* gene inducer. The wild type level of any given transcript is normalized to 1. Plots on the right show the relative transcription levels of *sco4699* and *sco4677* genes in the TopA-depleted and the wild type strains as well the control strain containing the integrative pIJ6902 vector. (**D**) Relative transcription of the *gyrB* gene and *sco4699* gene after exposure (0-60 minutes) to novobiocin at a final concentration of 10 µg/ml. The transcript enrichment for *gyrB* and sco4699 was normalized by the expression of the *hrdB* gene which served as the endogenous control and compared to the wild type strain in which transcription of analyzed genes was estimated as 1.

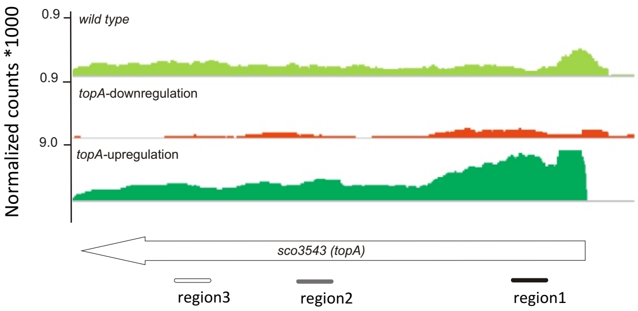


**Figure S5. The *topA* gene transcription and TopA binding.** RNA-Seq data visualization using Integrated Genome Viewer showing the number of successfully aligned fragments within the topA gene in the S. coelicolor wild type strain (light green), the TopA-depletion strain (orange) and the *topA*-induced strain (dark green). Note the Y-axis scale difference. The positions of the 1-3 regions tested in RT-qPCR (See also Fig. 4D) are marked

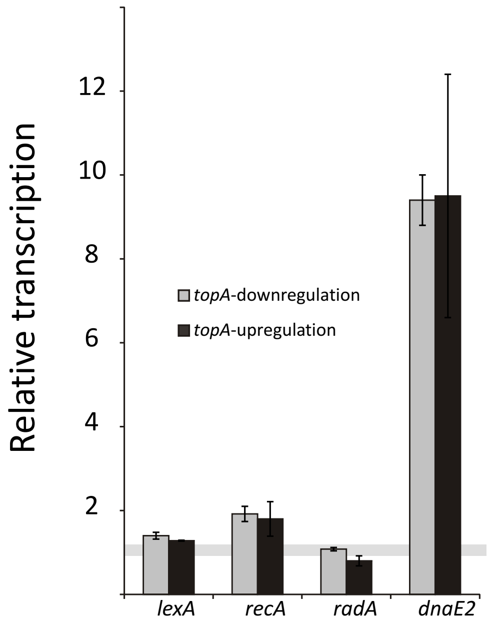


**Figure S6.** RT-qPCR analysis of the relative transcript level of genes belonging to the putative LexA regulon in *S. coelicolor (lexA, recA* and *dnaE)* under TopA-depletion (gray) and *topA*-induction (black) conditions. The *radA* gene, which does not belong to the *lexA* regulon, served as the negative control. The transcript enrichment for particular genes was normalized by the expression of the *hrdB* gene which served as the endogenous control and compared to the wild type strain in which transcription of analyzed genes was estimated as 1.
